# A rapid, inexpensive, culture-free, universal bacterial identification system using internal transcribed spacer targeting primers: a proof-of-principle study

**DOI:** 10.1101/2024.09.15.613074

**Authors:** Vishwaratn Asthana, Erika Martínez Nieves, Pallavi Bugga, Clara Smith, Tim Dunn, Satish Narayanasamy, Robert P. Dickson, J. Scott VanEpps

## Abstract

Techniques for bacterial detection and identification can be characterized as either untargeted and taxonomically broad or targeted and taxonomically narrow. Untargeted techniques (*e.g.*, culture and sequencing) are time-consuming and/or expensive while targeted techniques (polymerase chain reaction; PCR) can be faster and less expensive but require a strong pre-test suspicion of the potential organism to choose the correct test. We have developed a universal bacterial identification system that is as taxonomically broad as culture and amplicon sequencing but as fast, easy, and affordable as PCR. The platform utilizes a unique universal polymerase chain reaction (PCR) primer set that targets the internal transcribed spacer (ITS) regions between conserved bacterial genes, creating a distinguishable electrophoretic pattern for each bacterial species. Bioinformatic simulation demonstrates that at least 45 commonly isolated pathogenic species can be uniquely identified from a single set of PCR primers using this approach. We experimentally confirmed these predictions on seven representative human pathogens, including gram-negatives and gram-positives, aerobes and anaerobes, and spore formers. Without *a priori* knowledge of the organism, this system can rapidly identify the unique pattern generated by multiple species in a single reaction. Using quantitative PCR, the system can also determine the corresponding concentration of the organism in question. We also show that the primers are resilient to human DNA contamination at physiologic concentrations, eliminating the need for complex and time intensive extraction methods. Proof-of-principle testing on actual clinical specimens demonstrate that this assay can identify more than twice the species as current multiplex PCR assays (*e.g.*, BioFire) using only one universal primer pair in a single PCR reaction, providing results in <3 hours for <$20 without bioinformatic turnaround time.

**SIGNIFICANCE STATEMENT:** This paper describes a novel bacterial identification system that is as taxonomically broad as culture and amplicon sequencing but as fast, easy, and affordable as PCR. The assay covers more than twice the species as the leading multiplex PCR assays but uses only one universal primer pair in a single PCR reaction. It is resilient to human genomic contamination precluding the need for timely or costly methods to clear human DNA. This approach represents a major advancement in the decades-long struggle to rapidly and accurately identify bacteria.

## INTRODUCTION

Rapid, accurate detection and identification of bacterial species is a major challenge for human and veterinary clinical microbiology as well as food safety and environmental science. Current diagnostic techniques include culture with morphological/biochemical analyses and molecular assays such as polymerase chain reaction (PCR), next generation sequencing (NGS), mass spectroscopy, and immunodetection. These approaches can be characterized as either untargeted and taxonomically broad or targeted and taxonomically narrow in spectrum.

Untargeted techniques are those that do not require an *a priori* suspicion for a particular bacterial species and are therefore capable of detecting a broad – potentially hundreds – range of species. Culturing remains the most common untargeted technique and continues to be the gold standard for bacterial detection and identification. Although relatively cost-effective, it is time-consuming, often taking 12-72 hours to grow bacteria to the point where additional biochemical/morphological assays can be performed.[1] Furthermore, culturing can be limited by uncommon but relevant microorganisms that are fastidious or difficult to culture, requiring species-specific culture conditions to identify. Matrix-assisted laser desorption/ionization time-of-flight mass spectroscopy (MALDI-TOF) offers quick clinical turnaround on pathogen identity, but still requires pre-culture enrichment which suffers from the limitations noted above.[2] Untargeted molecular methods, such as metagenomic Next Generation Sequencing (mNGS), do not require culture but are expensive (can approach $1000/sample) and time consuming (1-5 days minimum) for both sample preparation and bioinformatic analysis, and are not widely available.[3-5]

Targeted techniques such as PCR and immunodetection are quick and inexpensive but species-specific and therefore require *a priori* knowledge of what the infectious organism might be in order to choose the correct test.[6, 7] Even a multiplex PCR format (*e.g.*, BioFire) does not cover a sufficient number of species (max 20 organisms) to replace culture as the gold standard for comprehensive detection/identification.[8] Specifically, when a test is limited to a subset of species, there is always a concern that a negative result only indicates that the causative organism was not included in the panel, rather than a true negative result. Furthermore, while an individual PCR test may be inexpensive, panels of dozens of targeted tests can approach thousands of dollars per sample.[7]

Here we describe a novel diagnostic approach that provides the speed, availability, utility, and cost effectiveness of PCR with the untargeted breadth of mNGS. This approach utilizes custom primers that target the regions between uniquely conserved regions in the bacterial genome, such as the 16s-23s internal transcribed spacer (ITS) region. While the 16s and 23s genes themselves are well conserved and show low variation between species, the ITS region between the two genes displays unique length and frequency heterogeneity that can be assessed using standard DNA fractionation methods (*e.g.*, electrophoresis; **Figure 1**).[9] A small number of additional ITS regions can be targeted and amplified to improve the specificity of the system. The use of these “universal” primers makes the system target-agnostic, meaning that the user does not need to have *a priori* knowledge of the bacteria in question when deciding what test or panel to use. In this study, we (1) describe the design of specific sets of universal primers capable of uniquely identifying up to 45 different clinically relevant pathogens *via* computational simulation; (2) experimentally verify the performance of those primers on a subset of those pathogens; (3) evaluate the potential for quantitative PCR (qPCR) to further enhance diagnostic utility; (4) confirm the specificity for bacterial vs human genomic DNA; (5) and demonstrate assay performance in clinically relevant matrices (*i.e.*, blood and urine). Taken together, these results provide a proof-of-concept for this approach to bacterial detection and identification.

**Figure 1:**
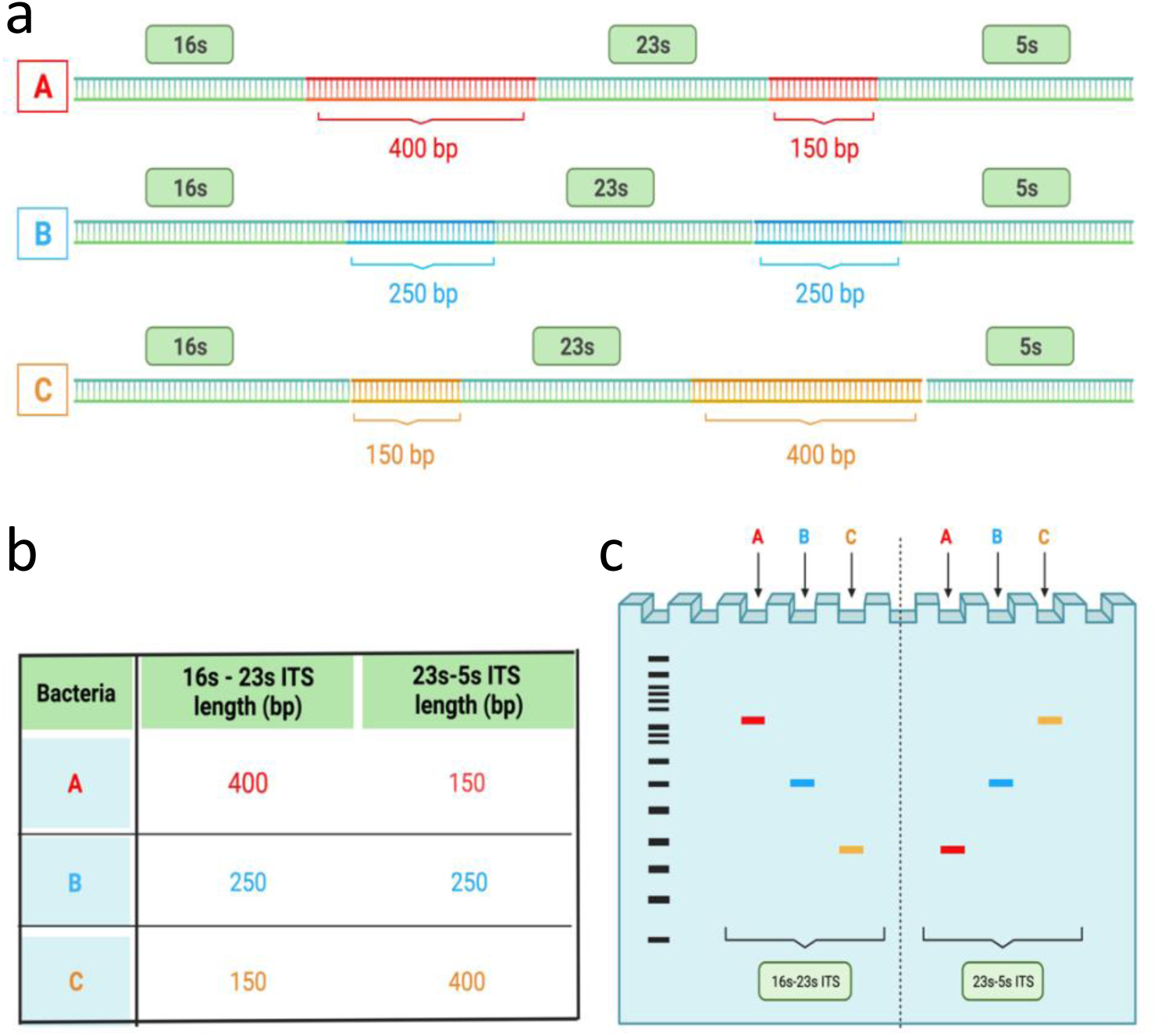
Conceptualization of the universal bacterial identification system. Universal primers targeting the flanks of conserved bacterial genes (including the 16s, 23s, and 5s ribosomal segments) can be used to PCR amplify the heterogeneous ITS regions producing a unique electrophoretic profile based on the length of the species-specific ITS regions. **(a)** Considering three hypothetical bacterial genomes (A, B, & C) with different ITS gap lengths, the resulting **(b)** amplicon lengths and **(c)** electrophoretic profiles demonstrate unique identifying signatures.

## METHODS

### Primer Design

First generation primers amplifying the 16s-23s ITS region were designed by scanning the 16s and 23s sequences for highly conserved regions. Second generation primers amplifying the 16s-23s and 23s-5s ITS regions were designed and modified based on prior literature.[10-12] Specifically, to improve the broad binding potential of these primers across diverse bacterial species, the universal primers were modified to include mixed bases (in a 1:1 ratio) as well as inosine, a universal base. In general, inosine was only included on the 3’ end of the primer to facilitate polymerase extension in the presence of a potential mismatch near the tail end of the primer binding site. Use of additional universal bases was avoided to prevent off-target binding and amplification. Final primer sequences can be found in **Supplementary Table 1**.

### Primer ITS amplification simulation

Following primer design, a Matlab® (R2021a) script was developed to establish primer binding sites and subsequent amplification profiles for each bacterial genome. In brief, the code searches each bacterial genome for primer binding sites by first probing for complementary base pair matches. This is followed by calculation of the binding affinity (ΔG) of each hit using integrated NUPACK^©^ code. Only primer binding sites with a ΔG < -9 kcal/mole were found to be adequately stable to form a dimer, a prerequisite for amplification. Next, each primer is elongated 5’ -> 3’ until it reaches the next inversely oriented primer binding site along the genome, the intervening sequence of which is considered an amplicon. Only those amplicons < 1,200 bp are included in the identity matrix generated for each universal ITS primer set. FASTA files containing bacterial genomic sequences were obtained from the National Center for Biotechnology Information (NCBI; see **Supplementary Table 2**).

### Cell culture and DNA extraction

Well-characterized laboratory strains of bacteria (**Supplementary Table 3**) were grown from single colonies overnight (12-18h) at 37°C and 200 rpm shaking in their respective medias (Lysogeny brothfor *Escherichia coli*, *Staphylococcus aureus, Klebsiella pneumoniae;* Muller Hilton media for *Pseudomonas aeruginosa;* Tryptic soy broth with 5% glucose for *Bacillus subtilis*). Cultures were then diluted and re-grown to an optical density at 600nm (OD_600_) of 0.4-0.6. DNA was extracted using a *Lucigen* purification kit according to the manufacturer’s instructions.

DNA was reconstituted in TE buffer and stored at -20°C. Purity and yields were quantified using a UV-Vis spectrophotometer (Thermo Scientific, Nanodrop 2000). Purified DNA was purchased for *Campylobacter jejuni* (ATCC 11168) and *Acinetobacter baumannii* (BAA-1605) from American Type Culture Collection (ATCC).

Clinical isolates of *E. coli* and *S. aureus* were detected, identified, and provided by the Michigan Medicine Clinical Microbiology Laboratory from positive blood or other sterile body fluid cultures obtained during usual patient care at Michigan Medicine. No patient identifying information was collected or viewed by the research team and therefore the study was deemed non-human research. For each species, 10 isolates from unique patient samples were utilized.

### Universal PCR amplification protocol

Bacterial DNA was amplified using iTaq polymerase from Bio-Rad according to the manufacturer’s instruction using the following thermocycling protocol: 1) 95°C for 3 min, 2) 95°C for 15s, 3) 60°C for 30s, 4) 72°C for 1min, and 5) Repeat steps #2 to #4 for 35 cycles. PCR reactants were subsequently separated *via* electrophoresis on a 10% polyacrylamide (PAGE) gel then stained with GelRed and imaged on a ChemiDoc XRS+ molecular imager (Bio-Rad). Gel images were analyzed using GelAnalyzer 19.1 (www.gelanalyzer.com). Specifically, the intensity as a function of distance from the well was plotted for each lane of the gel. Peaks were identified by comparison to standard 50 and 100 bp dsDNA ladders (New England Biolabs) that were fit to an exponential function of molecular weight versus distance.

### Quantification of bacterial concentration and assessment of limit of detection

*S. aureus* genomic DNA (150ng) was isolated and amplified as described above. It was then serially diluted 10-fold down to a final DNA yield of 3 fg. Each of these dilutions were then amplified for 60 cycles using SYBR green quantitative PCR (Bio-Rad CFX Opus 96 Real-Time PCR System) according to the kit instructions.

### Resilience to human genomic contamination

Human genomic DNA (Promega), isolated *E. coli* genomic DNA (as described above), or a combination of the two were amplified by PCR and analyzed by PAGE as described above. Three hundred seventy-five ng (2.5µLof 150 ng/µL) of human DNA and 75 ng (2.5µLof 30 ng/µL) of *E. coli* DNA were used as PCR templates. Reactions with increasing dilution ratios of *E. coli* to human DNA were created. Specifically, a 1:1 v:v ratio corresponds to 2.5 µL of 150 ng/µL (375ng) human genomic DNA and 2.5 µL of 30 ng/µL (75ng) *E. coli* genomic DNA, while 1:10 corresponds to 2.5 µL of 150 ng/µL (375ng) human genomic DNA and 2.5 µL of 3 ng/µL (7.5ng) *E. coli* genomic DNA, and so on. PCR reactions were run for either 30 or 60 cycles.

### Testing on contrived biofluid and real clinical samples

*E. coli* was grown overnight, diluted, then re-grown to mid-log phase as described above. Fifty µL of mid-log liquid culture was then added to 450 µL of porcine urine (graciously provided by the Mohammed Tiba Laboratory at the University of Michigan). This stock sample was then serially diluted 10-fold as above. Concentrations (CFU/mL) of the dilutions were confirmed by plating on LB agar followed by colony enumeration. DNA was subsequently extracted, PCR amplified, and analyzed by PAGE as outlined above.

Excess human blood from healthy volunteers, obtained from an IRB approved, informed consent exempt study (#HUM00143244) was collected in a NaCit Vacutainer tube. Then, 50µl of mid-log *E. coli* culture was added to 450ul of blood, and serial diluted as above. DNA was extracted, PCR amplified, and analyzed by PAGE.

In a pilot clinical study, three urine samples from patients with confirmed urinary tract infection (> 100,000 CFU/mL) were obtained from the Michigan Medicine Clinical Microbiology and Virology Laboratory via an IRB approved, informed consent exempt process (HUM00234440). Bacterial identity in these samples was confirmed via standard culture and MALDI-TOF by the Michigan Medicine Clinical Microbiology and Virology Laboratory (**Supplemental Table 4**). Five hundred µL of urine was centrifuged at 17,000 RCF for 10min. Supernatant was discarded, and the pellet was resuspended in Gibco DPBS 1x. The process was repeated twice, followed by DNA extraction, PCR amplification, and analysis by PAGE.

## RESULTS

We first set about testing whether our primers provide sufficient discrimination across common bacterial species using *in silico* analysis. Predicted amplification profiles of the 16s-23s ITS regions utilizing the 1^st^ generation universal primer set (**Supplementary Table 1)** were generated for each of 45 commonly isolated bacterial species (**Figure 2**). These data show a unique length profile for each species.

**Figure 2.**
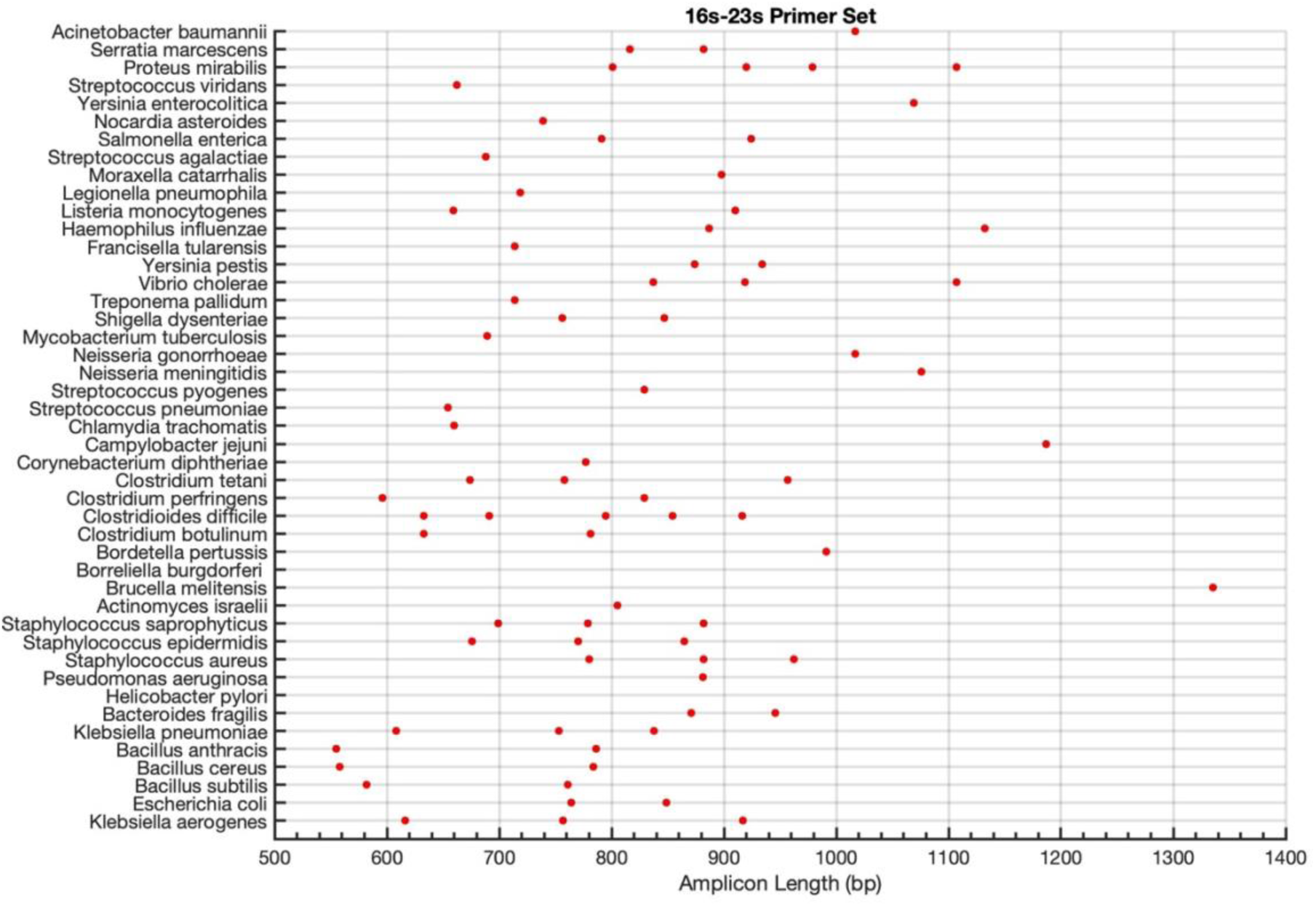
Simulated amplicon signatures. Expected length profile for 45 common clinical species using the 1^st^ generation 16s-23s universal primer set.

Having demonstrated the potential of the universal ITS primers *in silico*, we next ran proof-of-concept *in vitro* validation tests on DNA from seven representative bacterial species, including gram-negative and gram-positive bacteria, aerobes and anaerobes, and spore formers using the 1^st^ generation 16s-23s universal ITS primer set (**Figure 3**). These results demonstrate unique PCR product length profiles for each species that align with the simulated amplicon profiles (**Table 1**).

**Figure 3.**
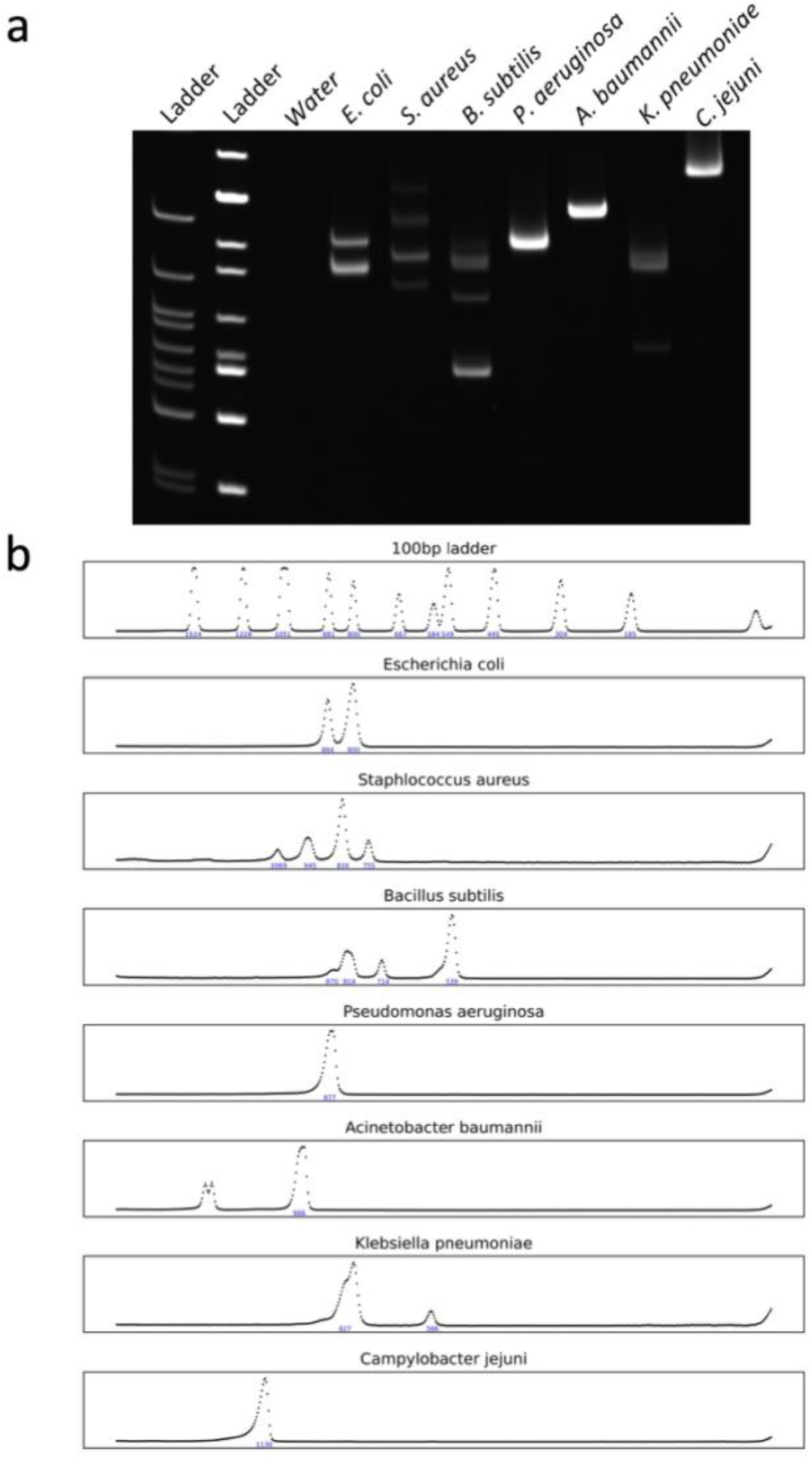
Experimental electrophoretic profiles generated using the 1^st^ generation universal 16s-23s ITS primers. **(a)** PAGE of PCR products from DNA isolated from different bacterial species and **(b)** image analysis results of the gel. Band analysis was restricted to PCR products < 1,200 bp which are unlikely to be spurious. Note the unique PCR amplicon profile for each species.

**Table 1:** Predicted vs observed PCR product lengths in bp and (number of associated repeats)

| Species | Predicted | Observed* |
| --- | --- | --- |
| *E. coli* | 764 (2), 849 (5) | 800±40, 884±44 |
| *S. aureus* | 780 (2), 882 (2), 962 (1) | 755±37, 834±41, 945±47, 1069±53 |
| *B. subtilis* | 582 (8), 761 (2) | 539±26, 714±35, 814±40 |
| *A. baumannii* | 1017 (6) | 986±49 |
| *P. aeruginosa* | 881 (4) | 877±43 |
| *K. pneumoniae* | 608 (1), 753 (4), 838 (3) | 586±29, 797±39, 827±41 |
| *C. jejuni* | 1187 (3) | 1130±56 |

\*Values include a 10% sizing error expected with gel electrophoresis

We next tested the ability to quantify the corresponding microbial concentration for an identified bacterial species as well as determine the limit of detection using 10-fold serial dilutions of *S. aureus* (**Figure 4**). The results confirm the log_2_-linear relationship between Cycle threshold (Ct) and concentration (Ct intervals of 3.3 per log_10_ dilution), even for the complex (more than one product) amplicons expected for *S. aureus*. Under the prescribed conditions, the system can detect as low as 30 fg of bacterial DNA per reaction, corresponding to 10 CFU with the limit of detection (LOD) limited primarily by the occurrence of primer dimers.

**Figure 4.**
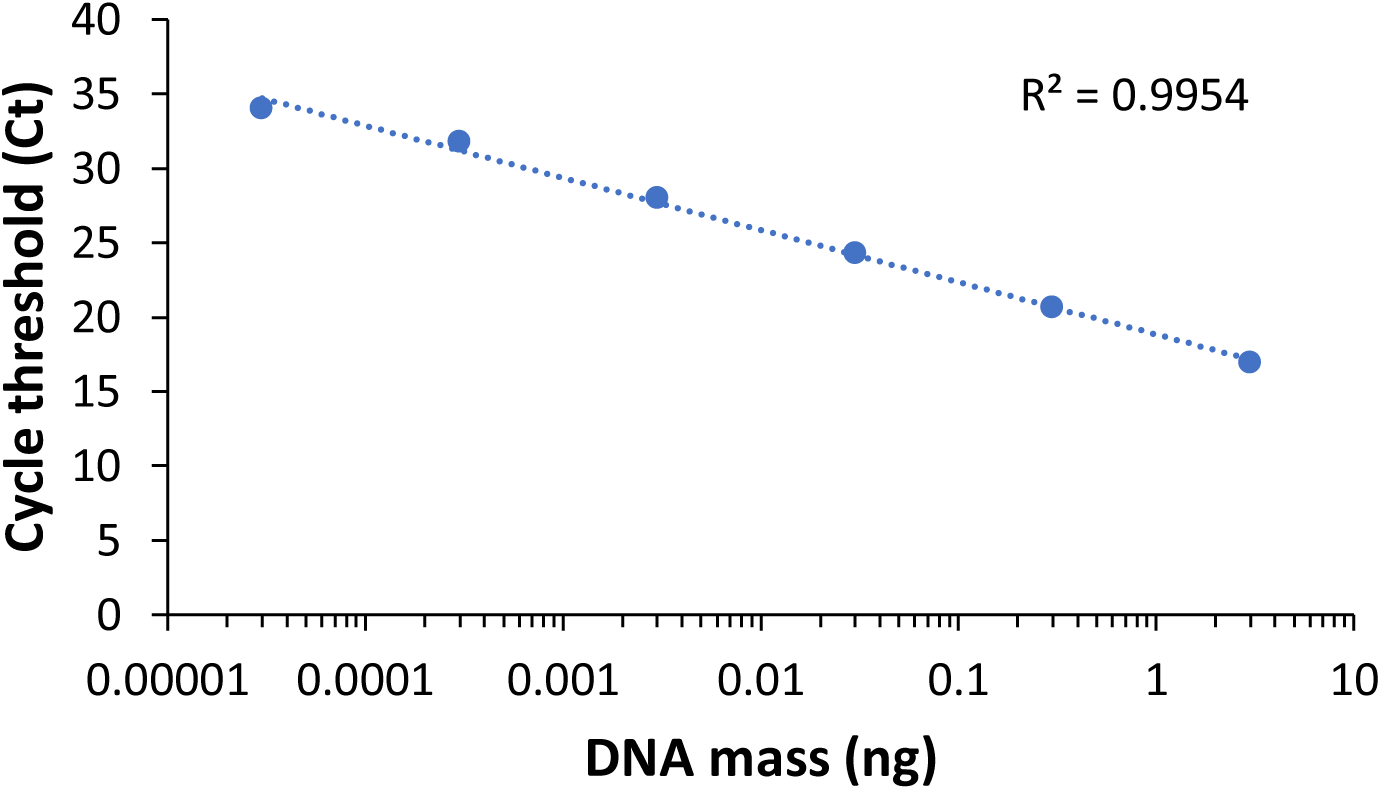
Quantification of bacterial DNA concentration and estimate of LOD using qPCR. *S. aureus* genomic DNA was serially diluted 10-fold and amplified using the 1^st^ generation universal 16s-23s ITS primers and SYBR green quantitative PCR. The results confirm the log_2_-linear relationship between PCR cycle and concentration (Ct intervals of 3.3 per log_10_ dilution) and demonstrate an LOD of 30 fg of bacterial DNA corresponding to 10 CFU.

Although this approach is as sensitive as any other PCR assay, its performance in the presence of large amounts of nontarget DNA (*e.g.*, human DNA) typically present in clinical samples remained unclear. Here we employed the 2^nd^ generation set of 16s-23s primers with improved specificity and breadth (see **Methods**). Specifically, the 1st generation of 16s-23s primers generated amplicons that were unnecessarily long, primarily because the primer binding sites were deeply embedded in the 16s and 23s rRNA genes, causing large portions of the resulting amplicon to be non-ITS, and therefore less discriminatory. Accordingly, we designed a 2^nd^ generation of 16s-23s primers with proximity to the 3’ and 5’ end of the 16s and 23s genes respectively. In addition, we added an inosine to the 3’ end of each primer to ensure robust amplification in the presence of potential 3’ mismatches at the primer binding site improving overall breadth. Lastly, the 2^nd^ generation 16s-23s primer set has less theoretical potential to bind non-specifically to the human genome as demonstrated by NCBI BLAST (data not shown). A second universal primer set targeting the 23s-5s ITS region was also generated. The simulated amplicon profiles (**Supplementary** Figure 1), number of ITS repeats (**Supplementary** Figure 2), and representative gel (**Supplementary** Figure 3) of these 2^nd^ generation primers demonstrate comparable performance; these primers were utilized henceforth for all subsequent experiments.

To mimic conditions that would be encountered in clinical specimens, where subject (*e.g.*, human) DNA can outnumber bacterial DNA by several orders of magnitude and potentially confound analysis, increasingly dilute concentrations of bacterial DNA were mixed with human genomic DNA and amplified (**Figure 5b**).[13, 14] The 2^nd^ generation 16s-23s universal primers do not amplify human genome within a typical number of PCR cycles (*i.e.*, 35 cycles). In addition, the presence of very high concentrations of human DNA does not interfere with the ability to amplify and detect bacterial DNA.

**Figure 5.**
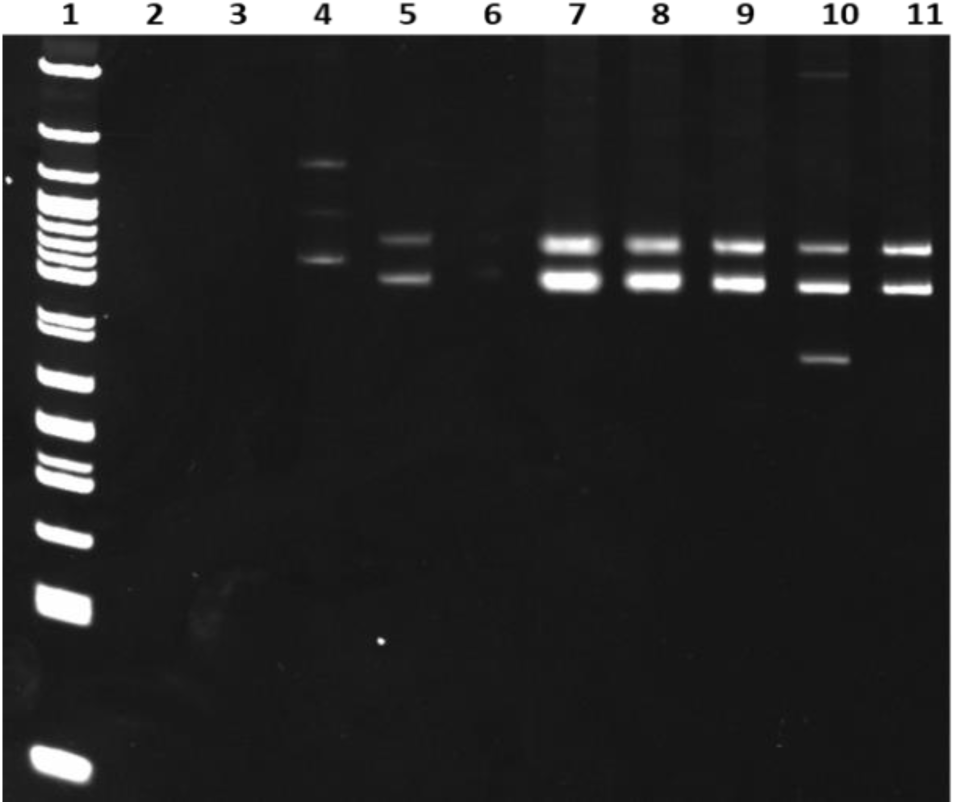
Resilience of the universal bacterial identification system to human genomic contamination. To assess specificity, human genomic DNA (375 ng), *E. coli* genomic DNA (75 ng), or a combination of the two were PCR amplified and visualized by PAGE. Lane 1: ladder; Lane 2: water; Lane 3: human DNA only, 35 cycles; Lane 4: human DNA only, 60 cycles; Lane 5: *E. coli* DNA only, 35 cycles; Lane 6: water; Lane 7: 1:1 human:*E. coli* DNA ratio, 60 cycles; Lane 8: 10:1 human:*E. coli* DNA ratio, 60 cycles; Lane 9: 100:1 human:*E. coli* DNA ratio 60 cycles; Lane 10: 1,000:1 human:*E. coli* DNA ratio, 60 cycles; Lane 11: 10,000:1 human:*E. coli* DNA ratio, 60 cycles.

Given that the system is resilient to off-target DNA, including human DNA, we next sought to determine whether the system could reliably detect multiple bacterial species in a single reaction (**Supplementary** Figure 4). We found that both *E. coli* and *B. subtilis* can be independently identified when present in near equimolar concentrations (*E. coli* and *B. subtilis* have similar genome sizes). However, amplification of the more dilute species appears to be hindered when present at less than a 1:10 concentration ratio by molarity relative to the dominant species.

To determine if this approach can be translated to clinical samples, where potential PCR inhibitors or other matrix constituents may confound analysis, we tested whether bacteria could reliably be detected and identified in two of the most commonly collected clinical biofluids – urine and blood – without culturing. Spiking serial 10-fold dilutions of *E. coli* into porcine urine had no effect on the expected PAGE band pattern (**Figure 6a**). In addition, when using the universal ITS primers in conjunction with SYBR green qPCR, the assay was able to reliably detect < 10^4^ CFU/ml in urine (**Figure 6b**). Similar PAGE results were obtained for serial dilutions of *E. coli* in whole blood (**Figure 6c**). Finally, we show that the system, in concordance with culture results, reliably identifies clinically relevant bacterial species in actual human urinary tract infection samples assayed in urine (**Figure 6d**). Specifically, sample 1 had a gel pattern consistent with *E. coli* while samples 2 and 3 had patterns consistent with *P. aeruginosa* (**Supplementary Table 4**).

**Figure 6.**
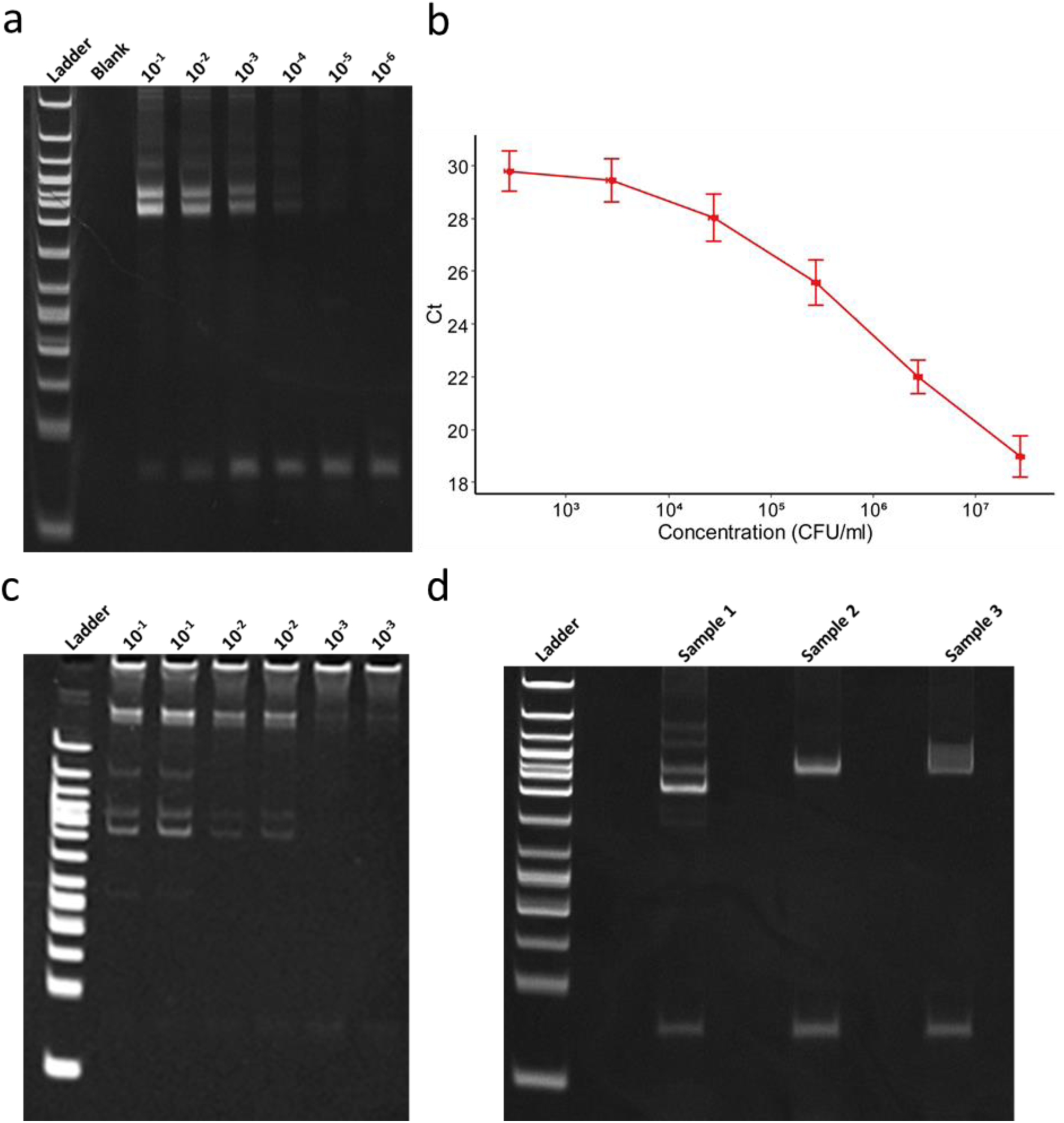
Assay performance in clinically realistic samples. **(a)** PAGE and **(b)** qPCR results of serial 10-fold dilutions of *E. coli* in porcine urine amplified using the universal primer set. Error bars represent standard error of 4 independent samples with triplicate measures. **(c)** PAGE results of serial 10-fold dilutions of *E. coli* in human blood. **(d)** PAGE results of human clinical urine samples with laboratory confirmed urinary tract infections (> 100,000 CFU/mL).

## DISCUSSION

In this proof-of-principle study, a single set of PCR primers was able to discern 45 bacterial species *in silico*. This was achieved using custom universal primers that bind to conserved bacterial genes and amplify intervening non-conserved gaps (*i.e.*, ITS). Because ITS regions are non-coding, and as a result not well conserved due to a lack of evolutionary pressure, these regions tend to have significant length and sequence heterogeneity between bacterial species.[15] This diversity in ITS length facilitates discrimination at the species level using simple experimental techniques (*i.e.*, electrophoresis). One limitation of our computational method is that we underestimated the ΔG binding affinity. This is due to the inability of commercially available nucleic acid thermodynamic software to incorporate the ΔG of mixed and/or universal bases. To be clear, the proposed approach is not theoretically limited to 45 species. Rather, the number of discernable species can increase based on the availability of bacterial species’ genome sequences. However, experimentally, gel electrophoresis can only resolve amplicons sized 10% apart, leading to some overlap between organisms in the current dataset.[16] To address this, additional universal primers targeting ITS regions between other conserved bacterial genes can be included to improve the specificity of the system by generating additional amplicon profiles. Indeed, this may eventually be required as closely related species, including those of the Klebsiella genus, generate PCR product length profiles with moderate overlap with the 16s-23s primer set alone. However, incorporation of the 23s-5s primer set generates additional length profiles that differentiates closely related species (**Supplementary** Figure 5). Another option would be to incorporate PCR product melt analysis to provide additional information with which to discriminate closely related species.[17] While not demonstrated here, the ratio of ITS repeats also has the potential to improve specificity when the reaction is not over cycled (*e.g.*, *B. subtilis* has two bands at an amplicon ratio of 1:4 while *K. pneumoniae* has three bands at an amplicon ratio of 1:4:3).

The ability to generate a unique length profile using universal ITS primers was experimentally validated in a cohort of seven species. While quite similar, experimental alignment to simulation was not perfect. We occasionally observed a spurious band that was not present in simulation. For example, *B. subtilis* demonstrates a heavier than expected band *via* PAGE that is not predicted by simulation. Concurrent 3% agarose gel analysis of the *B. subtilis* PCR products is missing this band, suggesting that that the band in question is unique to PAGE and likely the result of abnormal migration of a complex secondary structure (**Supplementary** Figure 6). This was confirmed by NGS analysis which demonstrated near complete alignment (98.2%) of all sequencing reads to the reference *B. subtilis* genome at the predicted target sites (data not shown). The use of PAGE for this proof-of-principle work provides a resolution limit for detecting PCR products. Grouping of amplicons predicted by simulation within a certain tolerance can be used to account for these resolution limits. For example, the predicted bands for *S. aureus* are 780, 875, 889, and 962 bp, which when grouped to within a 5% tolerance become 780, 882, and 962 bp bands. This can lead to some variation in band location and occasionally give the appearance of “missing” bands. Although the predicted amplicons do not perfectly align with experimental data, the spacing between bands is well preserved and the nonoverlapping readout predicted by simulation is maintained, the latter of which is essential for any future diagnostic. In addition, existing well described modalities such as capillary electrophoresis, which are both denaturing and have single base pair resolution, can potentially rectify this limitation.[18]

Ultimately, the final identity matrix that will be used to identify bacterial species can be constructed using simulation, experimental data, or both, which in turn will be used to generate a maximum likelihood estimate for a culprit organism. It should be noted that clinical isolates and strain level variations do not significantly alter simulation (**Supplementary** Figure 7) or experimental results (**Supplementary** Figure 8) using the current primer sets. It may be possible in the future to use additional primer combinations, targeting other conserved regions, to provide strain level differentiation while also maintaining species level identification, which may prove useful when identifying, for example, *E. coli* subspecies (Enterotoxigenic *E. coli* vs Enterohemorrhagic *E. coli*).

Without having *a priori* knowledge of the infectious organism, our proposed approach can also simultaneously determine the concentration of an identified microbe. Determining the concentration is clinically important. Providing a quantitative readout combined with clinical gestalt may help differentiate infectious from commensal bacteria and/or contaminants. For example, a pathogen load of < 10^5^ CFU/mL in urine is often considered non-infectious.[19] Concentration can be determined via incorporation of fluorescent qPCR which allows for automated real-time quantification (**Figure 4**). Of course, calibration against known concentration standards will be required to arrive at a colony count equivalent (CFU/mL).

PCR is theoretically capable of identifying even a single nucleic acid strand with sufficient cycling. Here we found that the LOD, using standard bacterial DNA extraction and amplification, was 10 CFU, which is well within the realm of clinical relevance for all major clinical compartments.[19-21] Methods of PCR amplifying bacterial DNA immediately from the biofluid of origin, as opposed to extracting the DNA first, could enhance sensitivity further.[22]

While both existing and emerging diagnostics, including matrix-assisted laser desorption/ionization time of flight mass spectroscopy (MALDI-TOF) and NGS, are sensitive in theory, they are limited by noise that originates from human cells and off-target genomic DNA. This is especially pronounced in blood where human cells and DNA can outnumber a bloodborne bacterial infection by several orders of magnitude.[13] The genomic targets of the custom primers described here, specifically the flanks of the 16s-23s and 23s-5s ITS regions, are specific to the bacterial kingdom such that off-target binding and amplification of the human genome is unfavorable and unlikely. Indeed, we show both *in silico* and *in vitro* that human DNA does not amplify within a conventional number of PCR cycles (*i.e.*, 35 cycles), and produces spurious bands when cycled at a non-conventional number (*i.e.*, 60 cycles). Even at the current LOD of the system, human genomic DNA does not interfere with bacterial DNA amplification and detection, despite being present up to 10^5^-fold in excess by mass. This potentially precludes the need for timely and costly human DNA extraction methodologies that have proved problematic for emerging bacterial diagnostic technologies.

Occasionally, clinical samples contain a mixture of pathogenic bacterial strains. In these infrequent scenarios, or situations where contaminant bacteria obscure the actual species of interest, the ability to discern multiple bacterial species in a single reaction is helpful. We theoretically show that multiple bacteria can simultaneously be detected when present within an order of magnitude of each other by molarity. Past this point, it appears the more concentrated species hinders amplification of less concentrated species, likely via consumption of PCR reagents and other mechanisms that have been explored in the literature.[23] For almost all applications, detecting the dominant and likely pathogenic species in a sample should be sufficient, however detecting additional species in solution is an area of further investigation. Of note, linear deconvolution could be used to break down the multiple length profiles from different bacteria into their constituent parts.

While turnaround time was not specifically tested, this system is theoretically capable of identifying a bacterial species within a few hours. Without optimization, for the approach described, DNA extraction takes approximately 1 hour, PCR requires 1 hour, and gel electrophoresis/imaging another 30 minutes for a total 2.5 hours. Eventually, the need for DNA extraction as performed here could be obviated using established methodologies that sequentially lyse and PCR amplify bacterial DNA in the biofluid of origin.[22] Thermocycling times could also be optimized, and gel electrophoresis/imaging simplified, the latter using established commercial capillary electrophoretic apparatuses, all of which could bring runtimes down to a goal time-to-result of under 1 hour.[24, 25]

Overall, this universal bacterial identification system represents a novel diagnostic approach to a longstanding clinical conundrum. There is currently no analogous technology that combines the performance of PCR with the target-agnostic benefits of NGS and culture. This is not a multiplex PCR platform that uses species-specific primers to assay an upper limit of 10 to 20 pathogens at a time.[6] Instead, it utilizes a parsimonious primer set to assay a much larger bacterial database, which could include hundreds of species, and is not limited to a single clinical compartment or bacterial classifier (e*.g.*, Gram-positive versus Gram-negative) like traditional bioassays.[26]

## AUTHOR CONTRIBUTIONS

VA, EMN, PB, RPD, and SV were responsible for designing experiments, analyzing data, and writing the manuscript. CS was responsible for designing and running experiments. TD and SN helped with analysis.

## ACKNOWELDGEMENTS

We would like to thank our collaborator, Dr. Tiba and his lab, for graciously providing urine samples. We would also like to thank Piyush Ranjan for his assistance with metagenomic analysis of our sequencing data.

## DATA AVAILABILITY

The datasets used and/or analyzed during the current study are available from the corresponding author on reasonable request.

## Supplemental Information for

**Supplementary Table 1:**
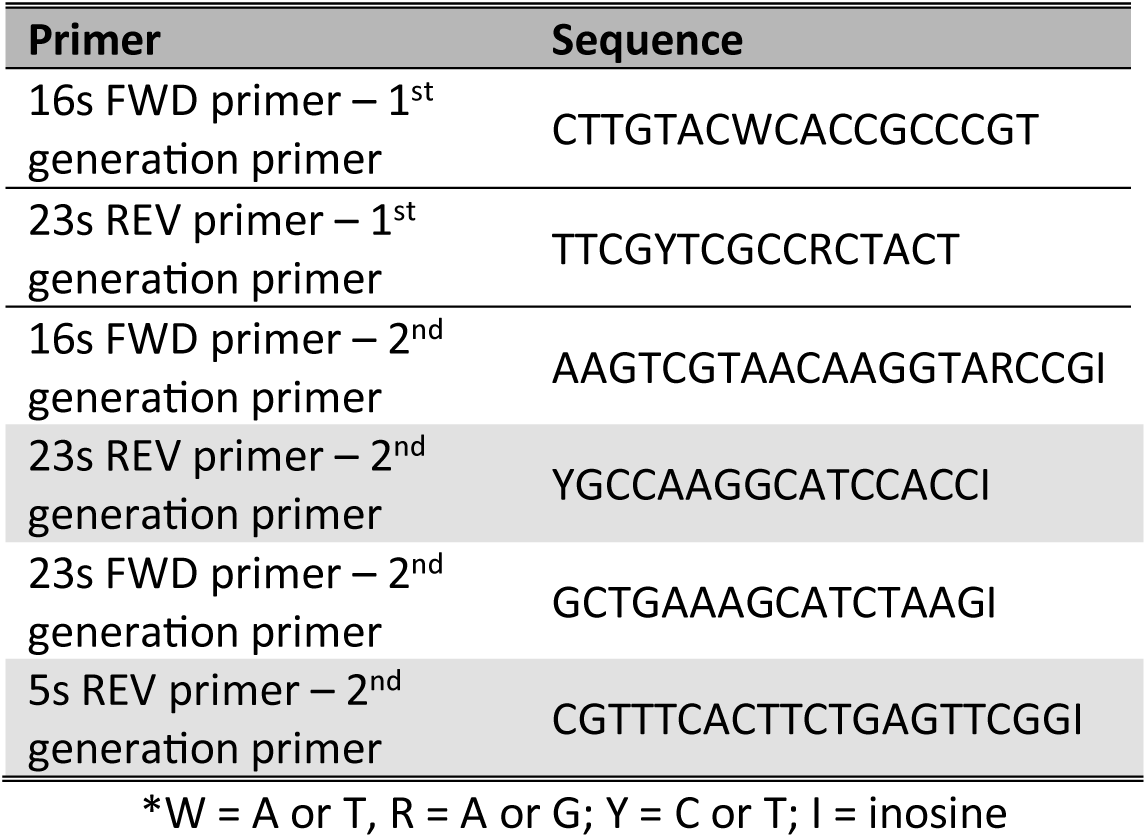
Universal ITS primer sequences*.

**Supplementary Table 2:** NCBI reference sequences.

| **Species** | **NCBI Reference Sequence** |
| --- | --- |
| *Klebsiella aerogenes* | NC_015663.1 |
| *Escherichia coli* | NC_000913.3 |
| *Bacillus subtilis* | NC_000964.3 |
| *Bacillus cereus* | NZ_CP034551.1 |
| *Bacillus anthracis* | NC_003997.3 |
| *Klebsiella pneumonia* | NC_016845.1 |
| *Bacteroides fragilis* | NC_006347.1 |
| *Helicobacter pylori* | NZ_LS483488.1 |
| *Pseudomonas aeruginosa* | NC_002516.2 |
| *Staphylococcus aureus* | NC_007795.1 |
| *Staphylococcus epidermidis* | NC_005003.1 |
| *Staphylococcus saprophyticus* | NZ_CP026337.1 |
| *Actinomyces israelii* | NZ_LR134357.1 |
| *Brucella melitensis* | NZ_CP026337.1 |
| *Borrelia burgdorferi* | NC_011728.1 |
| *Bordetella pertussis* | NC_018518.1 |
| *Clostridium botulinum* | NC_009495.1 |
| *Clostridioides difficile* | NC_009089.1 |
| *Clostridium perfringens* | NC_008261.1 |
| *Clostridium tetani* | NC_004557.1 |
| *Corynebacterium diphtheriae* | NZ_CP028841.1 |
| *Campylobacter jejuni* | NC_002163.1 |
| *Chlamydia trachomatis* | NC_010287.1 |
| *Streptococcus pneumoniae* | NC_003098.1 |
| *Streptococcus pyogenes* | NZ_CP028841.1 |
| *Neisseria meningitidis* | NZ_LR134525.1 |
| *Neisseria gonorrhoeae* | NC_002946.2 |
| *Mycobacterium tuberculosis* | NC_000962.3 |
| *Shigella dysenteriae* | NC_007606.1 |
| *Treponema pallidum* | NZ_CP003679.1 |
| *Vibrio cholerae* | NC_002505.1 |
| *Yersinia pestis* | NZ_CP033699.1 |
| *Francisella tularensis* | NZ_CP009633.1 |
| *Haemophilus influenzae* | NC_000901.1 |
| *Listeria monocytogenes* | NZ_LT906436.1 |
| *Legionella pneumophila* | NC_002942.5 |
| *Moraxella catarrhalis* | NC_014147.1 |
| *Streptococcus agalactiae* | NZ_CP021773.1 |
| *Salmonella enterica* | NC_003197.2 |
| *Nocardia asteroides* | NZ_LR134352.1 |
| *Yersinia enterocolitica* | NC_008800.1 |
| *Streptococcus viridians* | NZ_LR134266.1 |
| *Proteus mirabilis* | NZ_CP021694.1 |
| *Serratia mirabilis* | NZ_CP041233.1 |
| *Acinetobacter baumannii* | NZ_CP043953.1 |

**Supplementary Table 3:** Bacterial strains.

| Species | Strain |
| --- | --- |
| *Campylobacter jejuni* | ATCC 11168 |
| *Klebsiella pneumoniae* | ATCC 43816 |
| *Bacillus subtilis* | ATCC 6051 |
| *Staphylococcus aureus* | ATCC 35556 |
| *Escherichia coli* | ATCC 25922 |
| *Pseudomonas aeruginosa* | PA7 |
| *Acinetobacter baumannii* | BAA-1605 |

**Supplementary Figure 1.**
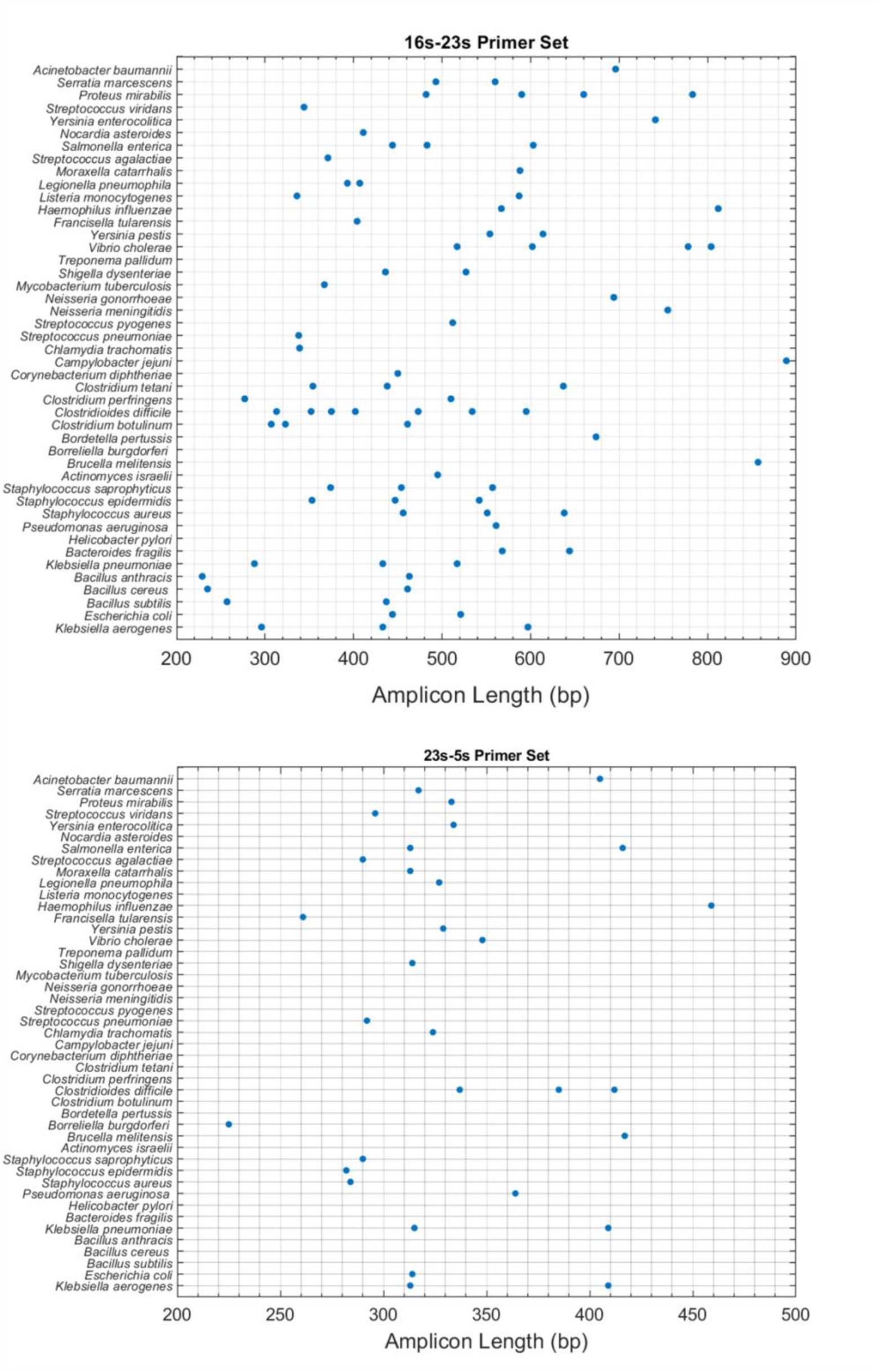
Simulated 2^nd^ generation universal primer amplicon signatures. Expected length profiles for 45 common clinical pathogens using the 2^nd^ generation **(a)** 16s-23s and **(b)** 23s-5s universal primer sets.

**Supplementary Figure 2.**
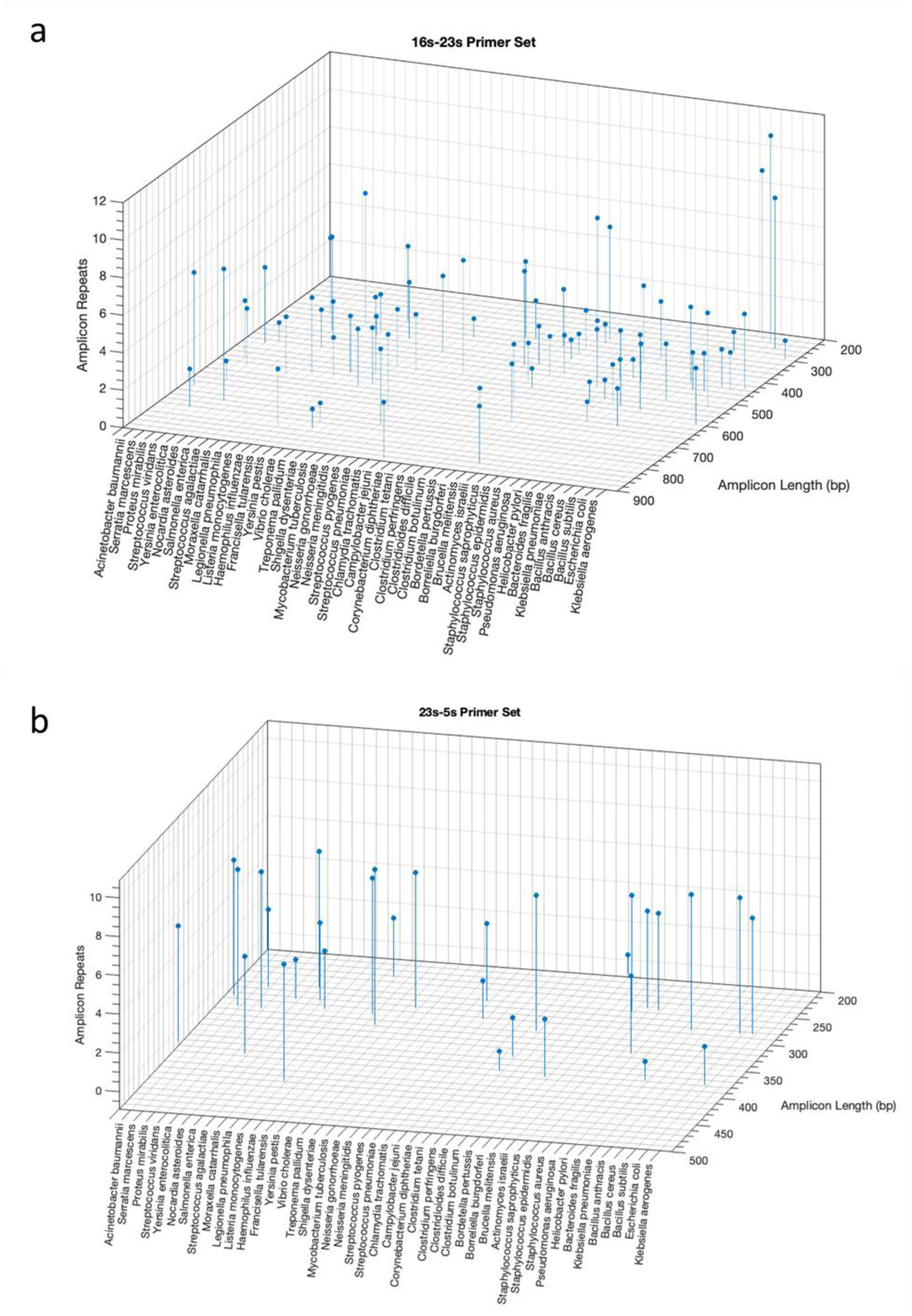
Simulated length profiles with associated repeats. Expected length profile and associated repeats for 45 common clinical pathogens using the 2^nd^ generation **(a)** 16s-23s and **(b)** 23s-5s universal primer sets.

**Supplementary Figure 3.**
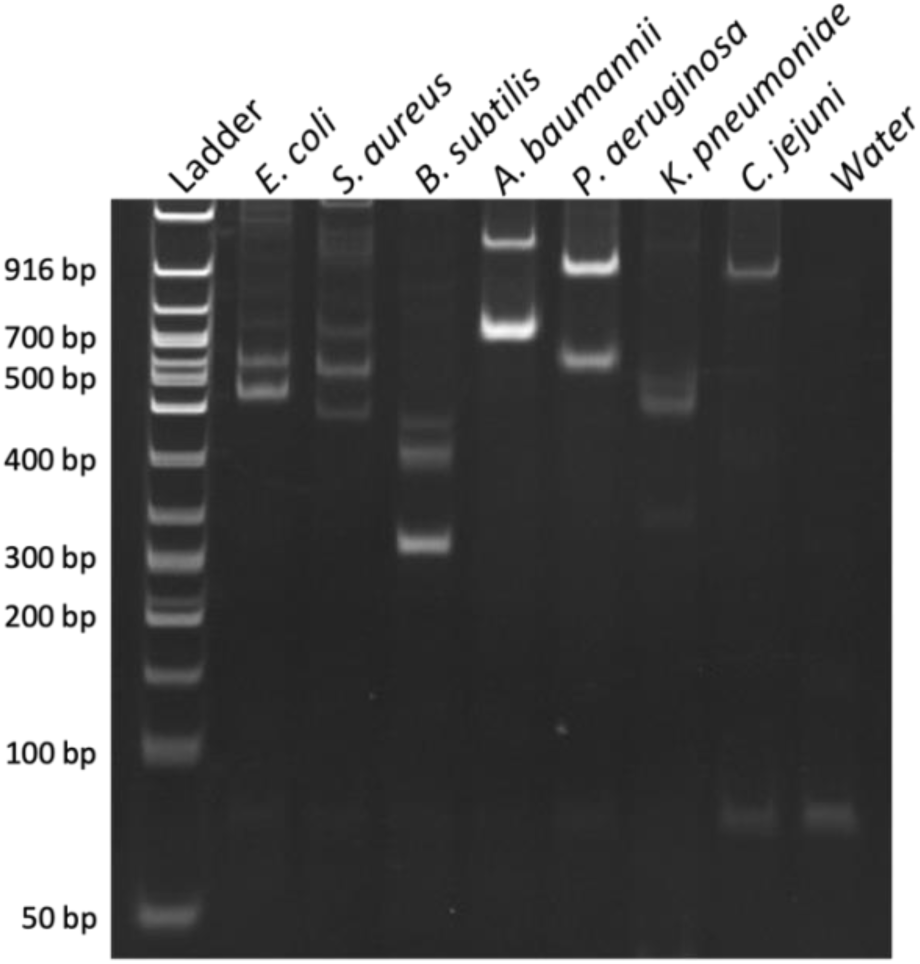
Experimental length profiles generated using the 2^nd^ generation universal 16s-23s ITS primers. PAGE of PCR products from DNA isolated from different bacterial species. The results mirror the bands generated by the 1^st^ generation universal 16s-23s primers, albeit lighter in weight.

**Supplementary Figure 4.**
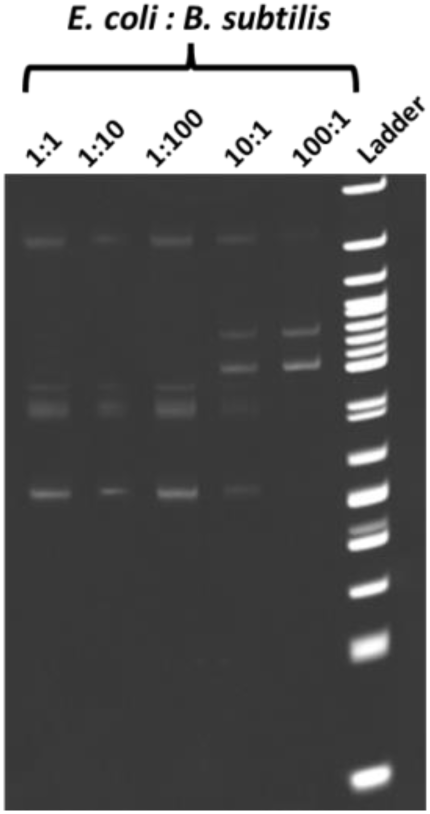
Amplifying various co-mixtures of bacteria in a single reaction. Varying ratios of *E. coli* to *B. subtilis* were PCR amplified for 35 cycles and visualized by PAGE. At a *E. coli* to *B. subtilis* ratio of 1:1, 75 ng of *E. coli* genomic DNA was mixed with 75 ng of *B. subtilis* genomic DNA. At 1:10, 7.5 ng of *E. coli* genomic DNA was mixed with 75 ng of *B. subtilis* DNA, and so on. Both species can be reliably identified when present within an order of magnitude of each other.

**Supplementary Table 4:** Clinical urine specimens.

| Sample | Culture identified species |
| --- | --- |
| Sample 1 | *E. coli* |
| Sample 2 | *P. aeruginosa* |
| Sample 3 | *P. aeruginosa* |

**Supplementary Figure 5.**
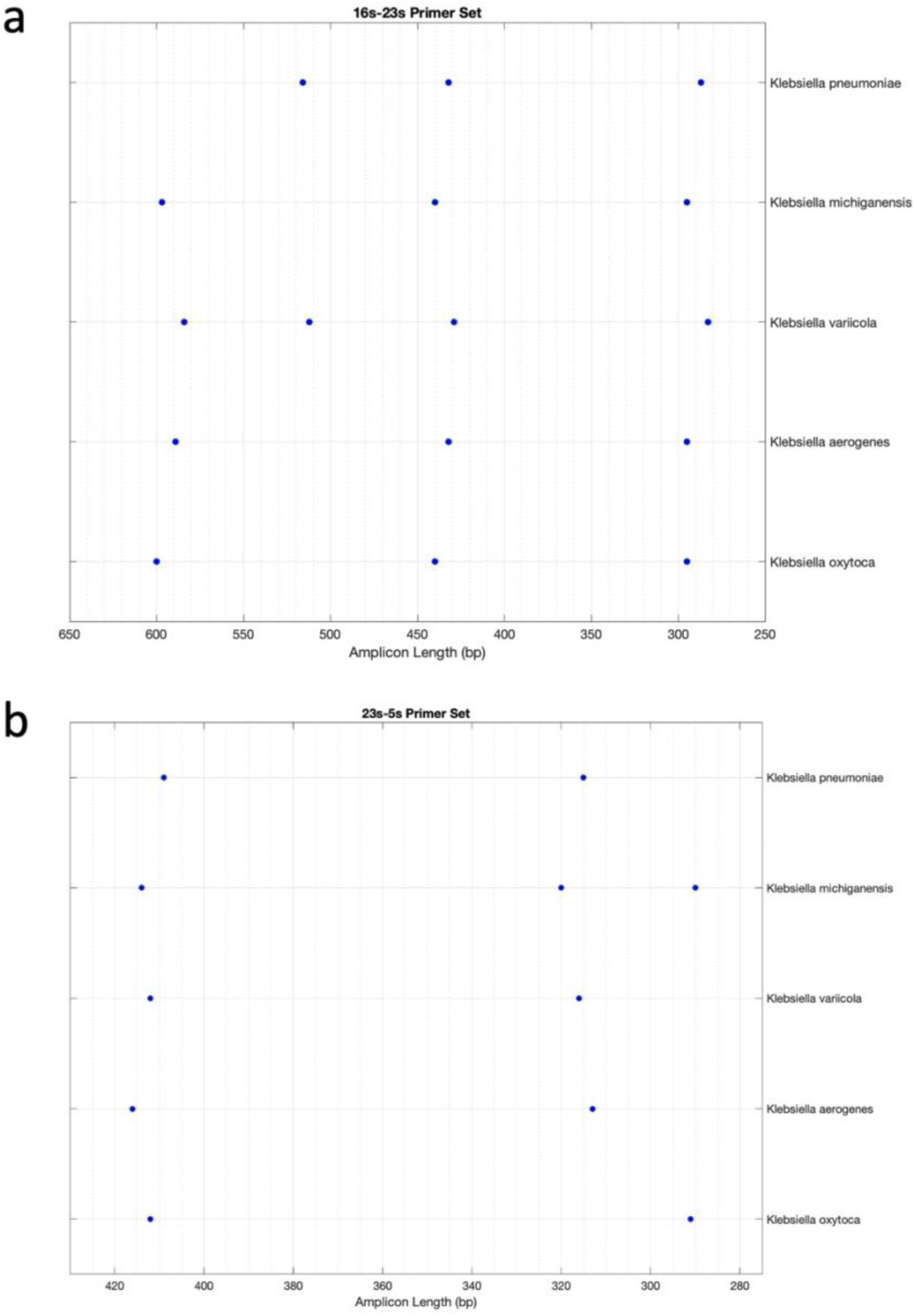
Simulated variation of closely related bacterial species. Expected length profile for five *Klebsiella* species using the **(a)** 16s-23s and **(b)** 23s-5s universal primer sets.

**Supplementary Figure 6.**
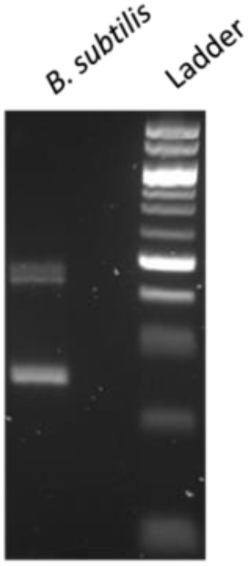
Resolving spurious bands using agarose gel electrophoresis. Post-PCR *B. subtilis* products were run on a 3% TAE agarose gel for 2.5 hours at 70 V. Only two bands are evident, instead of the three bands visible on PAGE, which is in line with simulation. Ladder is the 50 bp DNA ladder from NEB.

**Supplementary Figure 7.**
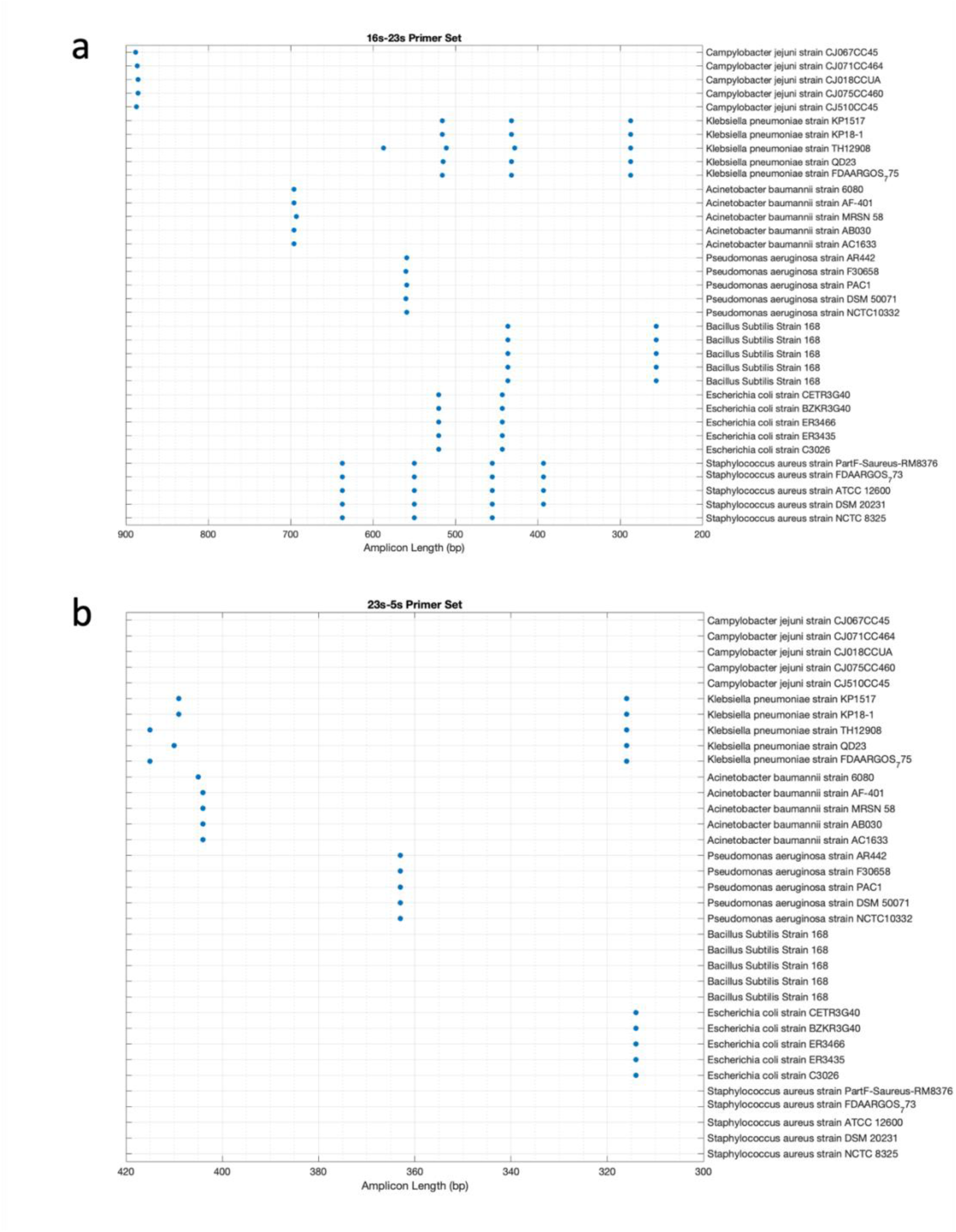
Simulated variation in strain/clinical isolate level length profiles. Length profiles for five strains/clinical isolates per bacteria for the seven bacteria experimentally tested in Figure 3 using the **(a)** 16s-23s and **(b)** 23s-5s universal primer sets.

**Supplementary Figure 8.**
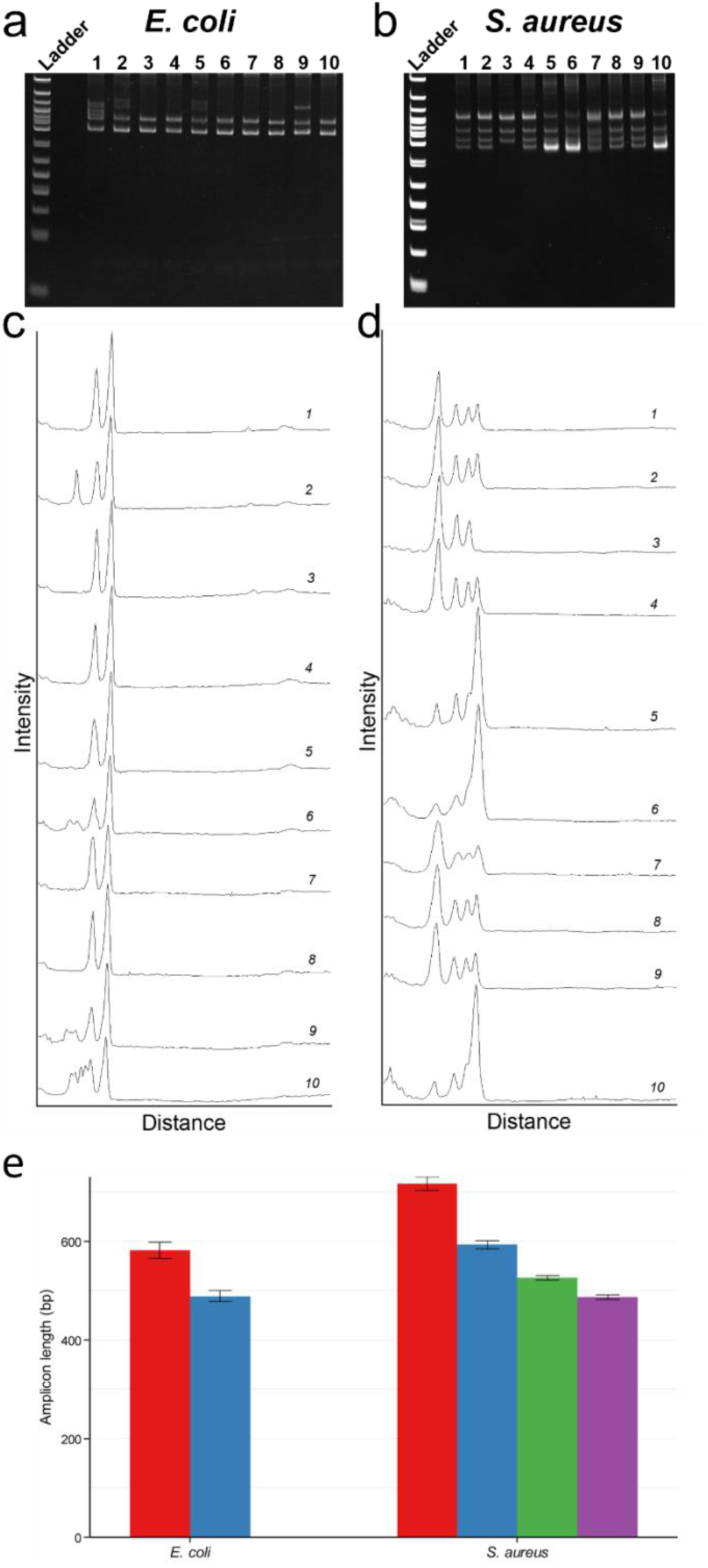
Experimentally validated variation in strain/clinical isolate level length profiles using the universal 16s-23s ITS primers. (a & b) PAGE of PCR products from DNA isolated from 10 different *E. coli* and *S. aureus* clinical isolates, (c & d) corresponding image analysis results of the gel, and (e) derived amplicon length mean and variation (error bars represent the standard deviation of 10 clinical isolates).

